# Phospho-proteomics reveals that RSK signaling is required for proliferation of natural killer cells stimulated with IL-2 or IL-15

**DOI:** 10.1101/2021.12.17.473192

**Authors:** Melanie A. MacMullan, Pin Wang, Nicholas A. Graham

## Abstract

Natural killer (NK) cells are cytotoxic lymphocytes that play a critical role in the innate immune system. Although cytokine signaling is crucial for the development, expansion, and cytotoxicity of NK cells, the signaling pathways stimulated by cytokines are not well understood. Here, we sought to compare the early signaling dynamics induced by the cytokines interleukin (IL)-2 and IL-15 using liquid chromatography-mass spectrometry (LC-MS)-based phospho-proteomics. Following stimulation of the immortalized NK cell line NK-92 with IL-2 or IL-15 for 5, 10, 15, or 30 minutes, we identified 8,692 phospho-peptides from 3,023 proteins. Comparing the kinetic profiles of 3,619 fully quantified phospho-peptides, we found that IL-2 and IL-15 induced highly similar signaling in NK-92 cells. Among the IL-2/IL-15-regulated phospho-sites were both well-known signaling events like the JAK/STAT pathway and novel signaling events with potential functional significance including LCP1 Ser5, PAK2 Ser141, and STK17B Ser12. Using bioinformatic approaches, we sought to identify kinases regulated by IL-2/IL-15 stimulation and found that the p90 ribosomal S6 kinase (p90RSK) family was activated by both cytokines. Using pharmacological inhibitors, we then discovered that RSK signaling is required for IL-2 and IL-15-induced proliferation in NK-92 cells. Taken together, our analysis represents the first phospho-proteomic characterization of cytokine signaling in NK cells and increases our understanding of how cytokine signaling regulates NK cell function.

## 1. Introduction

Natural killer (NK) cells are cytotoxic lymphocytes of the innate immune system that mediate anti-viral, anti-tumor, and other critical immune responses. The development, expansion, survival, and cytotoxicity of NK cells is regulated by many signals including cytokines, which are small, secreted proteins that bind to cell surface receptors and initiate intracellular signaling. Among the cytokines that regulate NK cells are the type I interleukins (IL) IL-2 and IL-15 (Abel et al., 2018). These two cytokines play distinct and complementary roles in promoting the growth and homeostasis of NK cells (de Rham et al., 2007; Mao et al., 2016; Pillet et al., 2009; Wu et al., 2017). However, the signaling pathways by which IL-2 and IL-15 control NK cell function are still not fully understood.

Despite low sequence homology, both IL-2 and IL-15 signal through heterotrimeric receptor complexes that include the IL-2/15Rβ subunit (CD122) and the common γ chain (γ_c_) (CD132) (Meghnem et al., 2017; Rodella et al., 2001). The third member of the receptor complex differs between the two cytokines, with IL-2 binding to complexes containing IL-2Rα (CD25), and IL-15 binding to complexes containing IL-15Rα (CD215) (Hamid et al., 2006; Tamzalit et al., 2014; Yang and Lundqvist, 2020). Upon cytokine binding, the receptor complex activates Janus family tyrosine kinases (JAKs) that phosphorylate and activate the signal transducer and activator of transcription (STAT) family of transcription factors as well as the Ras/Raf/MAPK and PI3K/AKT signaling pathways (Gotthardt et al., 2019; Marzec et al., 2008; Mishra et al., 2014; Mitra et al., 2015; Waldmann, 2002). Despite similarities in their shared receptor subunits and downstream signaling pathways, IL-2 and IL-15 can elicit different functions in immune cells. For example, mice deficient in IL-2 or IL-15 have distinct phenotypes, with IL-2 deficient mice suffering from overaccumulation of activated T and B cells while IL-15 deficient mice experience a lack of NK, NKT, and activated cytotoxic T cells (Ma et al., 2000). Further receptor knockout experiments in mice have demonstrated that IL-15/IL-15R signaling is essential for NK cell proliferation and development while IL-2/IL-2R is not (Dunne et al., 2001). Thus, there is great interest in understanding IL-2 and IL-15 signaling in immune cells.

Comparisons of IL-2 and IL-15 signaling have generated conflicting results. Although some studies have found that the cytokines produce indistinguishable signaling profiles (Zambricki et al., 2005), others have demonstrated substantial differences (Waldmann, 2015). Despite the functional relevance for NK cell function, studies of IL-2 and IL-15 signaling in NK cells have been limited to approaches such as immunoblotting (Frank et al., 1995; Gilmour et al., 2001) and phospho-flow cytometry (Pande et al., 2012) that can profile only a small number of known signaling events for which validated antibodies exist. In contrast, liquid chromatography-mass spectrometry (LC-MS)-based phospho-proteomics offers comprehensive, unbiased, and quantitative characterization of protein phosphorylation without the need for validated antibodies (MacMullan et al., 2019). As such, we sought to use LC-MS phospho-proteomics to compare the early signaling dynamics induced by these cytokines in the immortalized NK cell line NK-92 and then connect these signaling events to functional outcomes including proliferation (**Fig. 1**).

**Figure 1.**
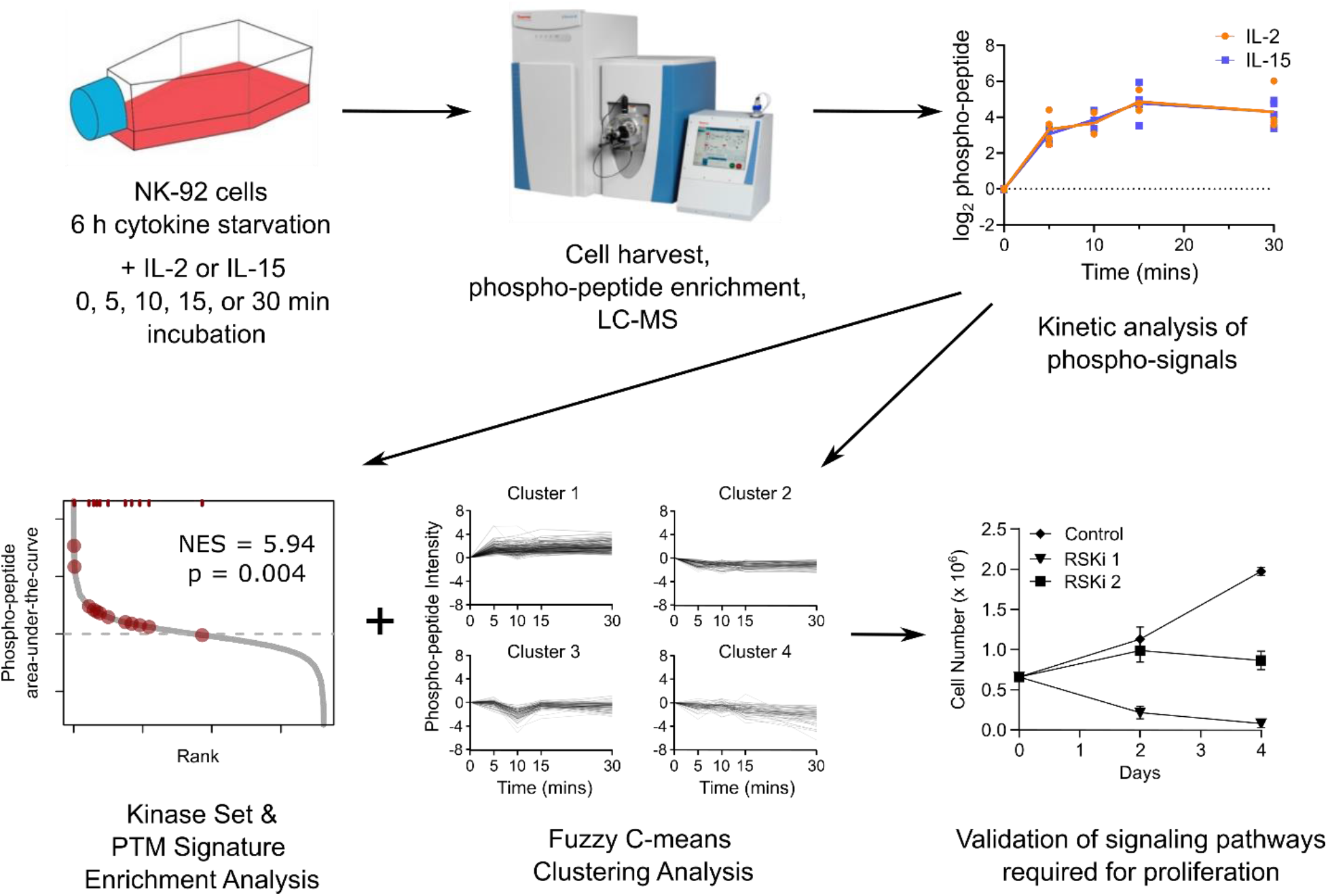
Experimental design for phospho-proteomic analysis of IL-2 and IL-15 signaling in NK cells. NK-92 cells were starved of cytokines for 6 h and then stimulated with 100 U/ml of either IL-2 or IL-15 for 0, 5, 10, 15, and 30 min. Cell lysates were harvested, trypsinized, and then enriched for phospho-peptides using TiO_2_ beads before analysis by LC-MS. The kinetic phospho-peptide data was analyzed using kinase set enrichment analysis (KSEA), post-translational modification signature enrichment analysis (PTM-SEA), and fuzzy C-means clustering. Lastly, based on the finding that both IL-2 and IL-15 induced RSK signaling, we used RSK inhibitors (RSKi) to validate that RSK signaling is required for proliferation of NK-92 cells stimulated with either IL-2 or IL-15.

## 2. Methods

### 2.1 Cell Culture

The immortalized NK-92 cell line was a gift from Dr. Jihane Khalife of the Children’s Hospital of Los Angeles. NK-92 cells were expanded in MEM Alpha (1x) + GlutaMAX media (Gibco 32561037) containing 8% fetal bovine serum (Sigma F2442), 8% horse serum (Gibco 16050130), 1% non-essential amino acids (Lonza BE13-114E), 0.2 mM myoinositol (Sigma I7508), 0.02 mM folic acid (Sigma F8758), 0.2 mM sodium pyruvate (Corning 25-000-Cl), 0.1 mM β-mercaptoethanol (Gibco 21985023), and 0.35% Pen Strep (Gibco 15140-122). Media was filtered using a 0.2 μm bottle filter and warmed to 37°C before being added to cells. Interleukin-2 (IL-2) (PeproTech 200-02) was added at a concentration of 100 U/mL during each media refresh. Cells were initially recovered at a concentration of 0.5 million cells/mL but maintained at a concentration of 0.25 million cells/mL for all subsequent passages. Cells were counted by trypan blue exclusion with a hemocytometer. Cells were grown in a 5% CO_2_, 37°C and humidified incubator and were used within 20 passages of thawing. For the experimental conditions, cells were starved for 6 hours in fresh media lacking cytokines and subsequently treated with 100 U/mL of either IL-2 or IL-15 (PeproTech 200-15).

### 2.2 Flow Cytometry Analysis

Following cell starvation, 0.5 million cells were treated with 100 U/mL IL-2 or IL-15 and 5 µg/mL Brefeldin A and allowed to incubate for 6 hours. Cells were then washed 1 time with 100 µL PBS and fixed and permeabilized in 100 µL Fixation and Permeabilizaton Solution from the BD Cytofix/Cytoperm Fixation/Permeabilization Kit (BD BioSciences 554714) for 10 minutes at 4°C. Cells were washed 2 times using 100 µL 1X Perm/Wash buffer provided in the Cytofix/Cytoperm Fixation/Permeabilization Kit and then labeled with a R-Phycoerythrin (PE) labeled anti-human interferon (IFN)-γ antibody at a ratio of 1:100 in 100 µL 1X Perm/Wash buffer and incubated for 30 min at 4°C. Cells were then washed 2 times using 100 µL 1X Perm/Wash buffer and resuspended in 100 µL PBS for flow cytometry analysis using a Miltenyi Biotec MACSQuant benchtop flow cytometer.

### 2.4 LC-MS Phospho-proteomics

Cell pellets were lysed in a modified urea-based lysis buffer (8 M urea, 100 mM Tris-HCl pH 7, 1 mM activated sodium vanadate, 2.5 mM sodium pyrophosphate, 1 mM β-glycerophosphate) by sonication in a cup-horn sonicator reservoir at 4°C. Protein concentration was measured using a BCA assay (Pierce P123227) and 1 mg of each sample was used for phospho-proteomics. Lysates were reduced with 5 mM DTT for 30 minutes at RT, alkylated with 25 mM iodacetamide for 30 minutes at RT in the dark, quenched with 10 mM DTT for 10 minutes at RT, and then acidified with 5% trifluoroacetic acid (TFA). Proteins were then digested to peptides using a 1:100 trypsin-to-lysate ratio by mass. Tryptic peptides were desalted by reverse phase C18 StageTips and eluted with 30% acetonitrile. The eluents were vacuum dried and then resuspended in loading buffer (80% acetonitrile (ACN), 6% TFA) to a concentration of 1 mg/mL for phospho-enrichment.

Titanium dioxide beads (PolyLC TT200C18) were weighed out at a ratio of 10 mg beads to 1 mg tryptic peptide for enrichment and resuspended in 30 µL per sample of suspension buffer (30% ACN, 0.1% TFA) to prepare a bead slurry. For each sample, 30 µL of bead slurry was aliquoted into a new tube. Reconstituted dried peptides were added to the bead aliquots and samples were rotated at RT for 30 min. A C8 StageTip was prepared and the bead-conjugated peptides were added. Non phospho-peptides were washed away and bead-bound phospho-peptides were eluted in two steps, first by 10% ammonium hydroxide and then by 80% ACN, 2% formic acid (FA). The eluents were vacuum dried and then desalted using C18 StageTips. Phospho-peptides were eluted with 50% ACN and eluents were vacuum dried. Dried phospho-peptides were resuspended in 6 µL of 0.1% FA.

The samples were randomized and 5 µL of each sample was injected onto an EASY-nLC 1200 ultra-high-performance liquid chromatography coupled to a Q Exactive Plus quadrupole-Orbitrap mass spectrometer (Thermo Fisher Scientific). Peptides were separated by a reverse phase analytical column (PepMap RSLC C18, 2 µm, 100 Å, 75 µm×25 cm). Flow rate was set to 300 nL/min at a gradient from 3% LC buffer B (0.1% formic acid, 80% acetonitrile) to 38% LC buffer B in 110 min, followed by a 10-min washing step to 85% LC buffer B. The maximum pressure was set to 1,180 bar, and column temperature was maintained at 50°C. Peptides separated by the column were ionized at 2.4 kV in positive ion mode. MS1 survey scans were acquired at the resolution of 70,000 from 350 to 1,800 m/z, with a maximum injection time of 100 ms and AGC target of 1e6. MS/MS fragmentation of the 14 most abundant ions were analyzed at a resolution of 17,500, AGC target 5e4, maximum injection time 65 ms, and normalized collision energy of 26. Dynamic exclusion was set to 30 sec, and ions with charge +1, +7 and >+7 were excluded.

MS/MS fragmentation spectra were searched with Proteome Discoverer SEQUEST (version 2.2, Thermo Scientific) against in silico tryptic digested Uniprot all-reviewed Homo sapiens database (release June 2017, 42,140 entries) plus all recombinant protein sequences used in this study. The maximum missed cleavages was set to two. Dynamic modifications were set to phosphorylation on serine, threonine or tyrosine (+79.966 Da), oxidation on methionine (+15.995 Da), and acetylation on protein N-terminus (+42.011 Da). Carbamidomethylation on cysteine (+57.021 Da) was set as a fixed modification. The maximum parental mass error was set to 10 ppm, and the MS/MS mass tolerance was set to 0.02 Da. The false discovery threshold was set strictly to 0.01 using the Percolator Node validated by q-value. The relative abundance of parental peptides was calculated by integration of the area under the curve of the MS1 peaks using the Minora LFQ node. Individual phospho-site localization probabilities were determined by the ptmRS node, and phospho-sites with <0.75 localization probability were removed. RAW LC-MS files and processed result files are available at MassIVE database under identifier MSV000088281 (reviewer login: MSV000088281_reviewer; reviewer password: a).

### 2.5 Data analysis and statistics

Proteome Discoverer peptide group abundance values of each sample were normalized to the sample’s median value. After normalization, samples were filtered for quantification in at least 50% of replicates in Perseus (Version 1.6.2.3) (Tyanova et al., 2016). The area under the curve (AUC) of each phospho-peptide time course was calculated using the composite trapezoidal rule (**Equation 1**):

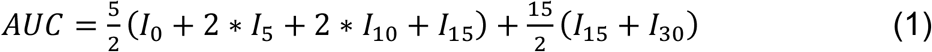

where *I*_*x*_ is the intensity of the phospho-peptide signal at *x* min. Kinase set enrichment analysis (KSEA) was performed using in-house Perl scripts and post-translation modification signature enrichment analysis (PTM-SEA) performed using the ssGSEA 2.0 R package available on Github (Krug et al., 2019). Fuzzy C-means clustering analysis was performed using the Mfuzz R package from Bioconductor in R version 4.0.2 (Kumar and Futschik, 2007). Additional data processing and analyses were performed in Microsoft Excel.

### 2.6 Drug Treatment

NK-92 cells were recovered in NK cell media for 24 hours before being counted by trypan blue exclusion and distributed into separate populations for drug treatment in triplicate. LJI308 (MedChemExpress, HY-19713) was used at a concentration of 20 µM, BI-D1870 (MedChemExpress, HY-10510) at a concentration of 2 µM and 10 µM, and DMSO was used as a control. Cell media and drug treatment were refreshed every 2 days, and cells were counted by trypan blue exclusion after each passage.

## 3. Results

### 3.1. IL-2 and IL-15 stimulate proliferation but not IFN-γ production in NK-92 cells

To understand how NK cells respond to the cytokines IL-2 and IL-15, we stimulated the immortalized NK cell line NK-92 with each cytokine individually. Notably, these cells are typically grown in media supplemented with IL-2. First, we tested the effects of these cytokines on proliferation and found that both cytokines induced robust cell growth (**Supp. Fig. 1A**). There was no significant difference in the growth rate of NK-92 cells stimulated with either cytokine. Next, we tested the effects of IL-2 and IL-15 on interferon (IFN)-gamma (γ) production and found that neither cytokine induced IFN-γ production (**Supp. Fig. 1B**). Taken together, this data demonstrates that both IL-2 and IL-15 induce proliferation but not IFN-γ production in NK-92 cells.

### 3.2. IL-2 and IL-15 activate canonical signaling pathways including JAK-STAT

Next, we sought to identify the signaling pathways induced by IL-2 and IL-15 using LC-MS-based phospho-proteomics. First, NK-92 cells were starved of cytokines for 6 h, and then cells were stimulated with either IL-2 or IL-15 for 0, 5, 10, 15 or 30 min (**Fig. 1**). Phospho-peptides were enriched using TiO_2_ beads and then identified and quantified using label-free LC-MS. In total, we quantified 8,692 phospho-sites on 3,023 proteins (**Supp. Table 1**). For subsequent quantitative analysis, we focused on the 3,619 phospho-peptides that were fully quantified across all 5 time points. To capture the total phospho-signal over time, we calculated the area under the curve (AUC) of the intensity for each phospho-peptide.

We then compared our phospho-proteomic data to the canonical IL-2 and IL-15 signaling pathways (**Fig. 2A-B**). One canonical signal induced by IL signaling is activation of JAK3 (Ikemizu et al., 2012; Morris et al., 2018), which is functionally bound to the *β*- and *γ*-chains of the IL-2/15 receptor (IL2/15R) along with JAK1. Our data revealed that both IL-2 and IL-15 robustly increased phosphorylation of JAK3 on Y785, an autophosphorylation site required for association with the adapter protein SH2B1 following IL-2 stimulation (Kurzer et al., 2004). We also quantified JAK3 peptides with phosphorylated residues S17 and S17/S20, although these phosphorylation sites had smaller AUC magnitudes than Y785 (average AUC values of 22.5 and 18.4 arbitrary units, respectively, compared to 47.8 arbitrary units for Y785) indicating less induction of phosphorylation over the 30-minute time period following stimulation.

**Figure 2.**
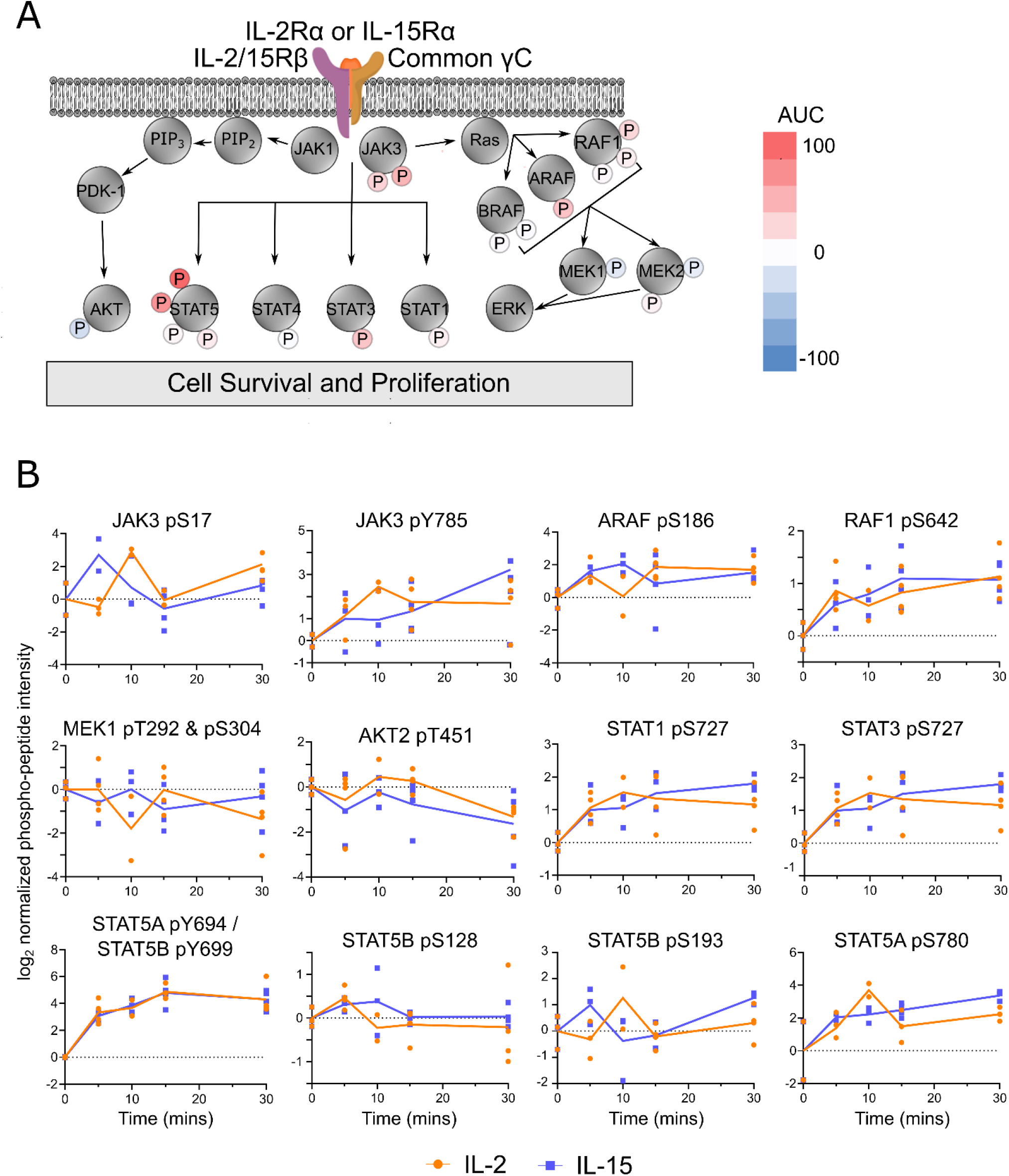
IL-2 and IL-15 activate canonical signaling pathways including JAK-STAT. A) Schematic of canonical signaling pathways downstream of the receptor complex of IL-2Rα, IL-2/IL-15Rβ, and the common γ chain. The small circles with P represent phosphorylation sites quantified by LC-MS phospho-proteomics. Each circle is colored from red to blue corresponding to positive and negative AUC values as shown. B) Kinetic profiles for selected phospho-peptides in canonical IL-2 and IL-15 signaling pathways. Each orange circle (IL-2) and blue square (IL-15) represent data from three biological replicates, and the line represents the average of the biological replicates.

One of the primary downstream targets of JAKs is the signal transducer and activator of transcription (STAT) family of transcription factors (Seif et al., 2017). Following IL-2/IL-15 stimulation, we observed increased phosphorylation on four phospho-sites on STAT family members including STAT1 S727, STAT3 S727, STAT5A Y694 / STAT5B Y699, and STAT5A S780. The phosphorylation of STAT5A Y694 / STAT5B Y699 (these tryptic phospho-peptides are indistinguishable) is crucial for activation and nuclear translocation of these transcription factors (Fahrenkamp et al., 2015; Klejman et al., 2002). Phosphorylation of S727 on both STAT1 and STAT3 has also been shown to be required for promotion of transcription activity under multiple cellular contexts (Decker and Kovarik, 2000). We also observed phosphorylation of STAT4A S733, STAT5B S128, and STAT5B S193, although the levels of these phospho-sites did not change in response to either IL-2 or IL-15 stimulation (i.e., AUC ≈ 0).

IL stimulation is also known to induce Ras/Raf/MAPK and PI3K/AKT signaling pathways (Marzec et al., 2008; Mishra et al., 2014; Mitra et al., 2015). Examination of phospho-signals from the Ras/Raf/MAPK pathway revealed one phosphorylation site on ARAF (S186), two phosphorylation sites on BRAF (S365 and S729), and three phosphorylation sites on RAF1 (S29, S259, and S642). Among these sites, only ARAF S186 and RAF1 S642 were significantly activated by IL-2 and IL-15 stimulation (average AUC of 38.5 and 24.9 arbitrary units, respectively). Downstream of RAF, we measured slightly decreased phosphorylation of the MAPK kinases MEK1 on T292/S304 and MEK2 on S23 and T394 in response to IL-2/15. In the PI3K/AKT pathway, we quantified only one phosphorylation site on AKT2 (pT451), and its phosphorylation was slightly decreased by IL-2 and IL-15 stimulation (average AUC of -17.1 arbitrary units). Taken together, these data support strong activation of the JAK/STAT signaling pathway and suggest weaker activation of the PI3K/AKT and Ras/RAF/MAPK signaling pathways in response to IL-2 and IL-15 in NK-92 cells.

### 3.3. IL-2 and IL-15 induce highly similar signaling in NK-92 cells

For all phospho-sites in the canonical signaling pathways, the kinetic profile of phosphorylation was qualitatively similar for IL-2 and IL-15-stimulated cells. We therefore tested whether this similarity between IL-2 and IL-15 signaling held true across all measured phospho-sites. Plotting the AUC values for all phospho-sites quantified in both IL-2 and IL-15-stimulated cells revealed a strong correlation (r = 0.72) (**Fig. 3A**). No phospho-peptides were significantly distant from the line of equality (i.e., where the AUC was equal for IL-2 and IL-15). For both cytokines, we observed a roughly equal balance of positive and negative AUC values (**Supp. Fig. 2**). Notably, by AUC, the most upregulated phosphorylation site in both IL-2 and IL-15-stimulated cells was STAT5A Y694 / STAT5B Y699 (average AUC of 115.3 arbitrary units). Interestingly, the serine/arginine repetitive matrix protein 2 (SRRM2) had both one of the most upregulated phospho-peptides (S1099/S1101/S1103, average AUC 97.1 arbitrary units) and one of the most downregulated phospho-peptides (S440, average AUC -58.5 arbitrary units) (**Fig. 3A-B**). We next searched the most up- and downregulated phospho-sites for sites with known regulatory function in the PhosphoSite Plus database (Hornbeck et al., 2015). Among the highly upregulated phospho-sites, this analysis highlighted plastin-2 (LCP1) S5 which regulates enzymatic activity and intracellular translocation (Koide et al., 2017) and the serine/threonine-protein kinase 17B (STK17B/DRAK2) S12 which negatively regulates IL-2 signaling and T cell development (Mandarano et al., 2020). Other phospho-sites with known regulatory function included vimentin (VIM) pS56 (Li et al., 2016; Ratnayake et al., 2021), the serine/threonine-protein kinase PAK 2 (PAK2) pS141 (Li et al., 2011; Zhan et al., 2003), Wiskott-Aldrich syndrome protein (WAS) pS483 (Cory et al., 2003; Liu et al., 2013), 40S ribosomal protein S6 (RPS6) pS236 & pS240 (Cerezo et al., 2021; Salizzato et al., 2016) and stathmin (STMN1) pS25 (Alesi et al., 2016; Kuang et al., 2016, 2015). Among the most downregulated phospho-sites with known regulatory function, we found stromal interaction molecule 1 (STIM1) pS608 (Casas-Rua et al., 2015; Pozo-Guisado et al., 2013; Tomas-Martin et al., 2015), mitochondrial fission factor (MFF) pS146 (Ducommun et al., 2015), and serine/threonine-protein phosphatase PP1γ catalytic subunit (PPP1CC) pT311 (Schmutz et al., 2011). Taken together, these data demonstrate that IL-2 and IL-15 induce highly similar signaling in NK-92 cells including many phospho-sites with known regulatory function.

**Figure 3.**
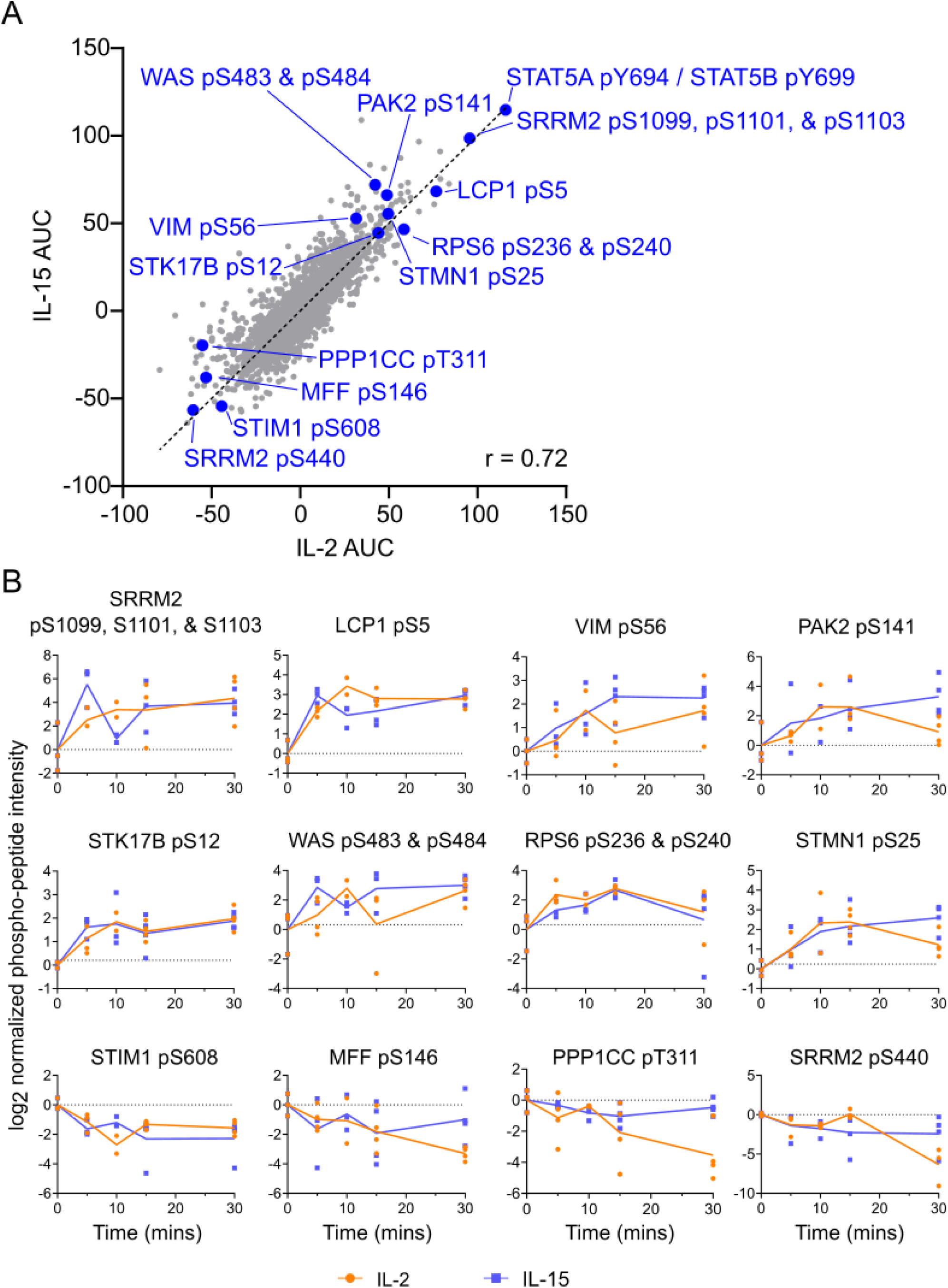
IL-2 and IL-15 induce quantitatively similar signaling in NK-92 cells. A) A scatter plot of AUC values reveals a strong correlation (r = 0.72) between IL-2 and IL-15 phospho-peptide time courses. Selected phospho-peptides with regulatory function according to the PhosphoSite Plus database (Hornbeck et al., 2015) are highlighted with blue circles. B) Kinetic profiles of selected phospho-peptides following IL-2 or IL-15 stimulation. Each orange circle (IL-2) and blue square (IL-15) represent data from three biological replicates, and the line represents the average of the biological replicates.

### 3.4. Enrichment analyses identify kinases activated by IL-2 and IL-15

Phospho-proteomics identifies the downstream phosphorylation sites but not the upstream kinases that perform the phosphorylation. We therefore used bioinformatic approaches to infer the kinases activated by IL-2 and IL-15 in NK-92 cells. First, we used post-translational modification signature enrichment analysis (PTM-SEA) (Krug et al., 2019) which tests for enrichment of kinase activity and signaling pathway signatures assembled from literature. Ranking IL-2 and IL-15-stimulated phospho-peptides by AUC, PTM-SEA identified p90RSK/RPS6KA1, PKCD/PRKCD, and RSK2/RPS6KA3 kinase signatures as significantly positively enriched in both IL-2 and IL-15 stimulated cells (**Fig. 4A**). Phospho-peptides from the IL-33 and EGFR1 signaling pathways were also significantly positively enriched in both IL-2 and IL-15 stimulated cells. Notably, although the signatures for p90RSK and RSK2 share four overlapping phospho-peptides (LCP1 pS5, RPS6 pS236, RANBP3 pS126, and BAD pS75), each kinase signature also had unique phospho-peptides including CARHSP1 pS52 and PDCD4 pS457 for p90RSK and STAT3 pS727 and TINF2 pS95 for RSK2 (**Fig. 4B**).

**Figure 4.**
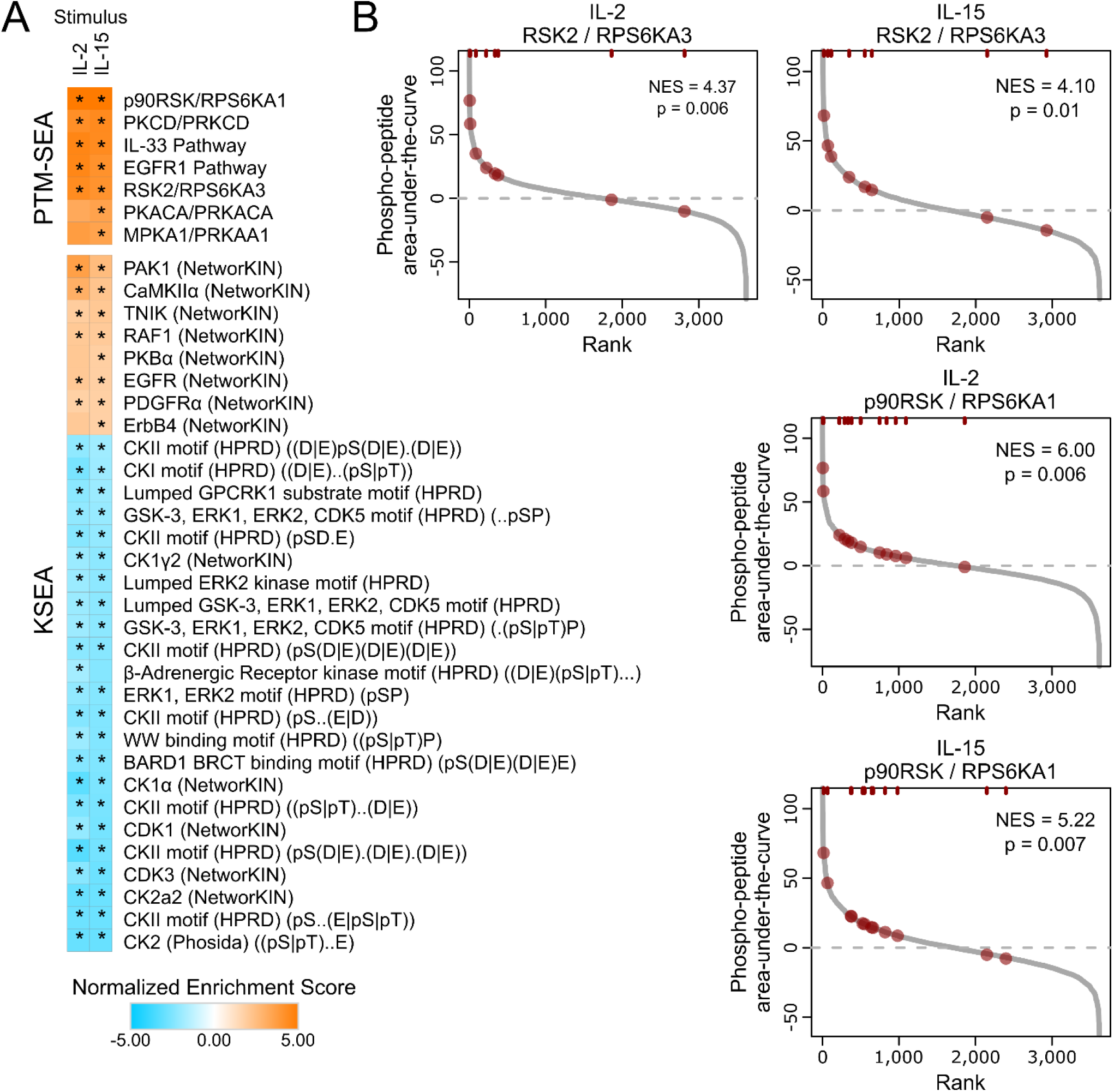
Post-translational modification signature enrichment analysis (PTM-SEA) reveals significant enrichment of p90RSK family kinases involved in propagating NK-92 cell signaling. A) PTM-SEA identified the p90RSK, RSK2, and the EGFR1 pathways as significantly (FDR < 0.01) upregulated in cells stimulated with either IL-2 or IL-15. Kinase set enrichment analysis (KSEA) corroborated significant upregulation of the RAF1 and EGFR kinases substrates and suggested downregulation of casein kinase (CK) II and GSK3/ERK substrates (FDR < 0.01). B) Rank-rank plots of both the RSK2 and p90RSK/RPS6KA1 kinase signatures reveal phospho-peptides identified in this dataset which overlap with those upregulated in the signature.

Next, we used kinase set enrichment analysis (KSEA), a technique that tests for statistical enrichment of kinase substrates predicted by NetworKIN (Linding et al., 2008) or phospho-peptides matching known kinase motifs (Keshava Prasad et al., 2009). In both IL-2 and IL-15-stimulated NK-92 cells, RAF substrates were significantly positively enriched, consistent with the known activation of Ras/RAF/MAPK signaling by IL-2/15 (Mishra et al., 2014; Nandagopal et al., 2014; Ross and Cantrell, 2018; Turner et al., 1993). In addition, EGFR substrates were enriched in both IL-2/15 stimulated cells, consistent with results from PTM-SEA. Among the most downregulated motifs were multiple casein kinase (CK) II motifs (e.g., pSD.E where. represents any amino acid) or GSK-3/ERK motifs (e.g., [pS|pT]P). Notably, nearly all kinase substrate sets enriched in IL-2-stimulated cells were also enriched in IL-15-stimulated cells. Taken together, the PTM-SEA and KSEA results suggest that IL-2 and IL-15 activate similar kinases in NK-92 cells including the p90RSK family of kinases.

### 3.5. Fuzzy C-means clustering identifies differing phospho-signaling kinetic profiles

Because our analysis of AUC compared phospho-peptides by their magnitude of signaling but not their kinetics, we next sought to identify different temporal patterns of phospho-peptide signaling downstream of IL-2 and IL-15. Using the most up- and downregulated phospho-peptides (i.e., top 10% by absolute value of AUC, n=724), we performed fuzzy C-means clustering analysis (Kumar and Futschik, 2007), a soft clustering approach which identifies common signaling trajectories while allowing phospho-peptides to belong to multiple clusters if appropriate (Hu et al., 2015; Rahmatbakhsh et al., 2021; Yang et al., 2015). This analysis revealed six distinct clusters with different kinetic profiles (**Fig. 5**). Cluster 1 demonstrated rapid and sustained upregulation of phosphorylation, whereas Cluster 2 showed rapid and sustained decreased phosphorylation. Interestingly, both Clusters 1 and 2 contained primarily IL-15 phospho-signals (74.0% and 65.9%, respectively). Cluster 3 and Cluster 5 also showed opposite kinetic profiles with initially decreased and increased phosphorylation, respectively, followed by adaptation to basal levels. Finally, Cluster 4 demonstrated slow but sustained decreases in phosphorylation, whereas Cluster 6 represented minimal initial signaling followed by increased phosphorylation at 10 and 15 min after stimulation with IL-2 or IL-15. Interestingly, phospho-peptides stimulated by both IL-2 and IL-15 which belong to the p90RSK signature (e.g., RPS6 pS236 & pS240) and the EGFR1 pathway signature (e.g., STAT3 pS727) were identified in Cluster 1. This analysis further suggests that both p90RSKs and the EGFR pathway are rapidly activated and sustained over 30-minute period following IL-2/15 stimulation in NK-92 cells.

**Figure 5.**
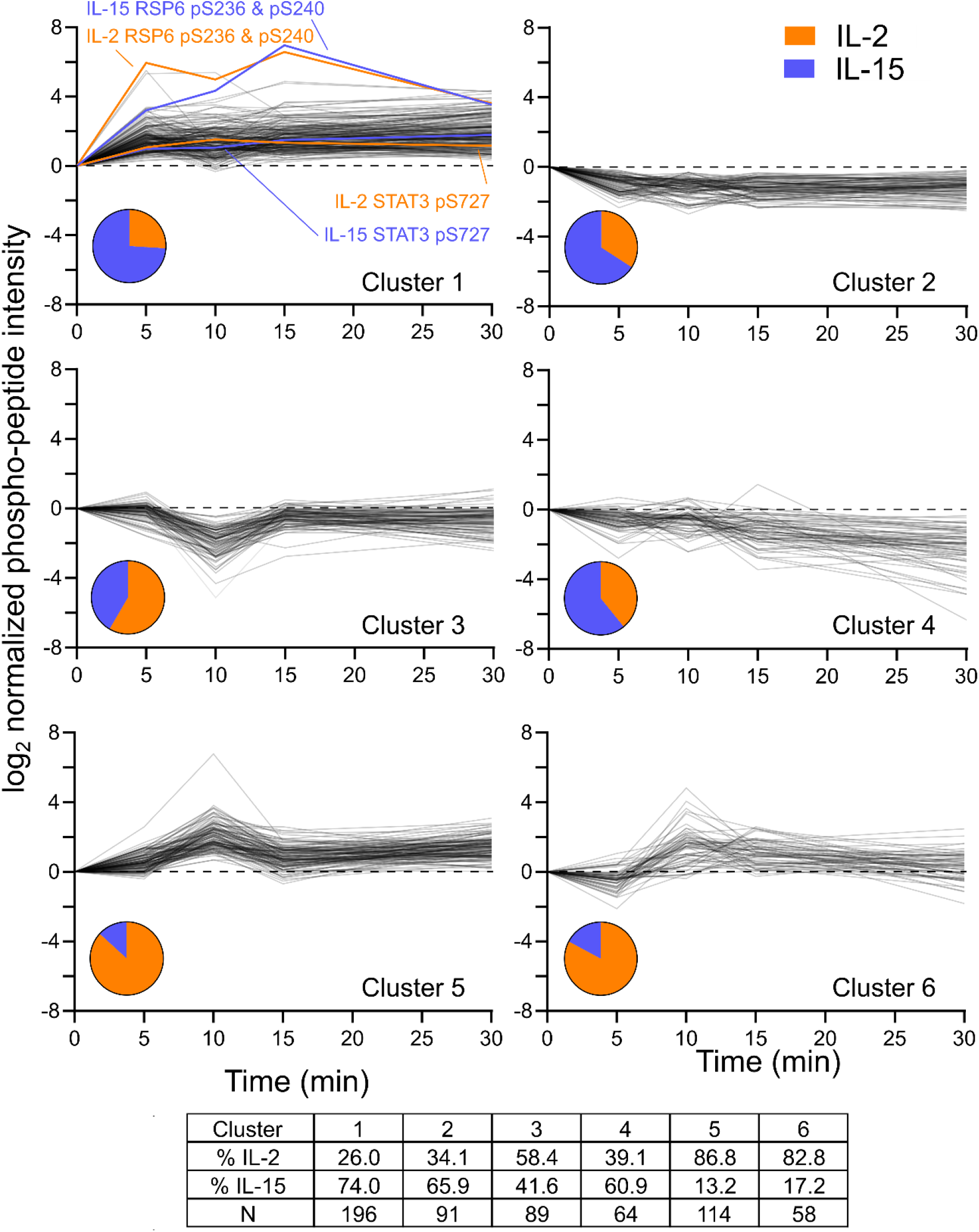
Fuzzy C-means clustering reveals distinct kinetic profiles of signaling downstream of IL-2 and IL-15. A) The top 10% most up- and downregulated phospho-peptides by absolute value of AUC (n=724) were subjected to fuzzy C-means clustering. 516 phospho-peptide trajectories were assigned into 6 distinct clusters (i.e., membership score > 0.7). Phospho-peptides corresponding to p90RSK (RSP6 pS236 & pS240) and EGFR1 pathways (STAT3 pS727) fell into cluster 1. The table shows the number and distribution of IL-2 and IL-15 stimulated phospho-peptides that were assigned to each cluster.

### 3.6. p90RSK signaling is required for IL-2- and IL-15-mediated proliferation

Following identification that p90RSK and RSK2 signaling was upregulated in IL-2 and IL-15-stimulated NK-92 cells, we sought to test whether p90RSK signaling is required for proliferation induced by IL-2 and IL-15. We therefore tested whether inhibiting RSK signaling with two structurally distinct p90RSK inhibitors, LJI308 and BI-D1870, would inhibit proliferation of NK-92 cells. Indeed, both inhibitors significantly reduced the growth of NK-92 cells treated with IL-2 or IL-15 (**Fig. 6**). This data demonstrates that p90RSK signaling is required for IL-2 and IL-15-mediated proliferation in NK-92 cells.

**Figure 6.**
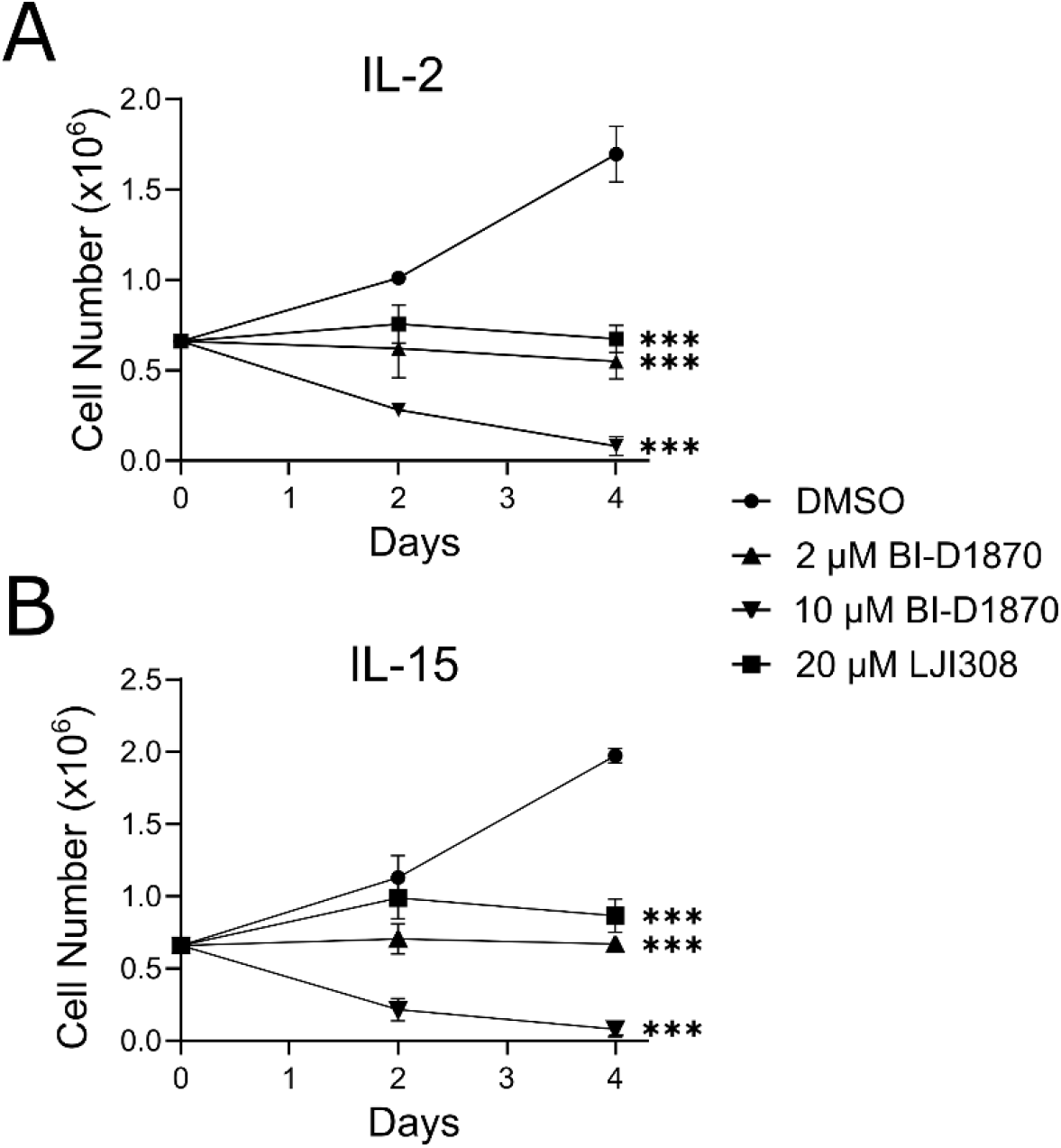
p90RSK signaling is required for IL-2- and IL-15-mediated proliferation of NK-92 cells. NK-92 cells were treated with the RSK inhibitors LJI308 (20µM), BI-D1870 (2 and 10 µM), or DMSO control, and cell growth was monitored for 4 days. Both RSK inhibitors significantly reduced cell growth compared to the DMSO control (***, Student’s t-test p-value < 0.001).

## 4. Discussion

Although it is widely recognized that interleukins including IL-2 and IL-15 regulate NK cell function, the signaling pathways stimulated by these cytokines are still not fully understood. To address this deficit, we present here the first LC-MS phospho-proteomic characterization of IL-2 and IL-15 signaling in the immortalized NK cell line NK-92. By quantitative analysis of the early signaling kinetics, we show that IL-2 and IL-15 activate highly similar intracellular signaling in NK-92 cells, including both known (e.g., JAK/STAT (Gotthardt et al., 2019; Morris et al., 2018)) and novel (e.g., p90RSK) signaling pathways. Remarkably, despite the low homology of these cytokines, the magnitude of signaling in response to IL-2 and IL-15 was extremely high across >3,600 phosphorylation sites (r=0.72) (**Fig. 3A**). Finally, using pharmacological inhibitors, we then demonstrated that p90RSK signaling is required for IL-2/15-mediated proliferation of NK-92 cells. Taken together, our results show the power of LC-MS phospho-proteomics to elucidate NK cell function and outline a pipeline for the analysis of signaling kinetics and downstream function in immune cells.

Previous studies of IL-2 and IL-15 signaling in NK cells were limited in that they require validated, costly antibodies to detect phospho-signals (Frank et al., 1995; Gilmour et al., 2001; Pande et al., 2012). In contrast, our LC-MS phospho-proteomic approach enables an unbiased and quantitative characterization of phospho-signaling across thousands of phosphorylation sites. Prior to our analysis, proteomic approaches to analyze NK cell signaling have been limited to a kinase-selective phospho-proteomic approach that identified and quantified 313 phosphorylation sites on 109 kinases at one time point (König et al., 2012). In contrast, our study identified >8,000 phosphorylation sites and fully quantified >3,600 phosphorylation sites across five time points. Thus, this work has greatly extended the landscape of known signaling events in NK cells. Among the phospho-sites we quantified downstream of IL-2 and IL-15, the most upregulated phosphorylation site was STAT5A Y694 / STAT5B Y699 (these tryptic phospho-peptides are indistinguishable). Remarkably, in terms of AUC, this phosphorylation site was 20% more upregulated than even the second most upregulated phospho-peptide (SRRM2 S1099/S1101/S1103). Notably, phosphorylation of these tyrosine residues on is crucial for activation and nuclear translocation of STAT5A / STAT5B (Fahrenkamp et al., 2015; Klejman et al., 2002). We also observed increased phosphorylation on STAT1 S727 and STAT3 S727, both of which are required for promotion of transcriptional activity under multiple cellular contexts (Decker and Kovarik, 2000). Taken together, this data demonstrates that IL-2 and IL-15 significantly upregulate the JAK/STAT pathway within 30 minutes of stimulation in NK-92 cells.

Our analysis also revealed that IL-2/15 activate the p90RSK family of kinases in NK-92 cells. Previous work has shown that IL-2 and IL-15 can activate p90RSK signaling in T cells (Franklin et al., 1996; Lin et al., 2008), but we are unaware of previous reports showing this connection in NK cells. Notably, our phospho-proteomic data showed only marginally increased phosphorylation of p90RSK (RPS6KA1) on S380, a phosphorylation site often used as marker of p90RSK activation (AUC values of 8.4 and 4.1 arbitrary units for IL-2 stimulation and IL-15, respectively) (Salhi et al., 2015; Singh et al., 2019). However, our bioinformatic analysis (e.g., PTM-SEA) revealed the activation of p90RSK from the enrichment of known downstream phosphorylation targets including the 40S ribosomal protein S6 (RPS6) S236, a downstream target of p90RSK and RSK2 that induces enzymatic activity and cell growth (Lin et al., 2019; Peng et al., 2010; Roux et al., 2003). In addition, the phosphorylation of plastin-2 (LCP1) S5 and serine/threonine-protein kinase 17B (STK17B/DRAK2) S12, both targets of p90RSK and RSK2 (Machado et al., 2021; Schenk et al., 2017), was significantly upregulated in response to IL-2 and IL-15. Both LCP1 S5 and STK17B S12 have previously been linked to IL-2 signaling in cytotoxic CD8+ T cells (Ross et al., 2016). Other notable p90RSK targets in our data set include RANBP3 pS126 (Yoon et al., 2008) and BAD pS75 (Cronin et al., 2021).

While p90RSKs have been shown to be rapidly activated by IL-2, IL-15, and IL-7 in T cells (Lin et al., 2008), their role in NK cells has not yet been elucidated. Previous research has suggested that CREB activation might be a result of RSK-2 activation in NK cells stimulated by IL-2, but this has not yet been conclusively demonstrated (Ponti et al., 2002). Importantly, when we treated NK-92 cells with the RSK inhibitors BI-D1870 and LJI308, we observed a complete suppression of cell growth and even cell death at higher doses, thus connecting our phospho-proteomic findings to NK cell function. Future work will seek to identify whether single or multiple members of the p90RSK family are required for proliferation of NK-92 cells.

In summary, this work presents quantitative insight into NK-92 cell signaling downstream of IL-2 or IL-15 and offers the potential for improved understanding of NK cell biology and immunoengineering. NK cells rely on a balance between activating and inhibitory receptors to initiate activity, making it challenging to characterize the activation signals that can be used for engineering NK cells for therapeutic usage (Paul and Lal, 2017; Sivori et al., 2019). Because of their complexity, NK cells are typically engineered with tools that have been optimized for T cells, resulting in suboptimal engineered cell designs (Albinger et al., 2021). Regardless, given that the immortalized cell line NK-92 is being used clinically for tumor immunotherapy (Cheng et al., 2012; Tang et al., 2018), our results may provide insight into the essential components of IL-2/15 signaling to improve engineering of NK-92 cells for optimal activity *in vivo*. As the field of immunotherapy continues to expand and to push into the clinic, phospho-proteomic characterization of immune cell signaling will be critical to the improved understanding of the complex mechanisms by which immune cell function is regulated and how immune cells can be manipulated for therapeutic benefit.

## CRediT Author Statement

**Melanie A. MacMullan:** Conceptualization, Methodology, Software, Validation, Formal analysis, Investigation, Data Curation, Writing – Original Draft, Writing – Review & Editing, Visualization, Project Administration; **Pin Wang:** Conceptualization, Resources, Writing – Review & Editing, Supervision, Funding Acquisition; **Nicholas A. Graham:** Conceptualization, Software, Resources, Writing – Review & Editing, Visualization, Supervision, Project Administration, Funding Acquisition.

## Abbreviations

AUC: area under the curve
IFN: interferon
IL: interleukin
JAK: Janus family tyrosine kinase (JAKs)
KSEA: kinase set enrichment analysis
LC: liquid chromatography
MS: mass spectrometry
NES: normalized enrichment score
NK: natural killer
PBS: phosphate buffered saline
PTM-SEA: post-translational modification signature enrichment analysis
RT: room temperature
STAT: signal transducer and activator of transcription

## FIGURES AND FIGURE LEGENDS

**Supplemental Figure 1.**
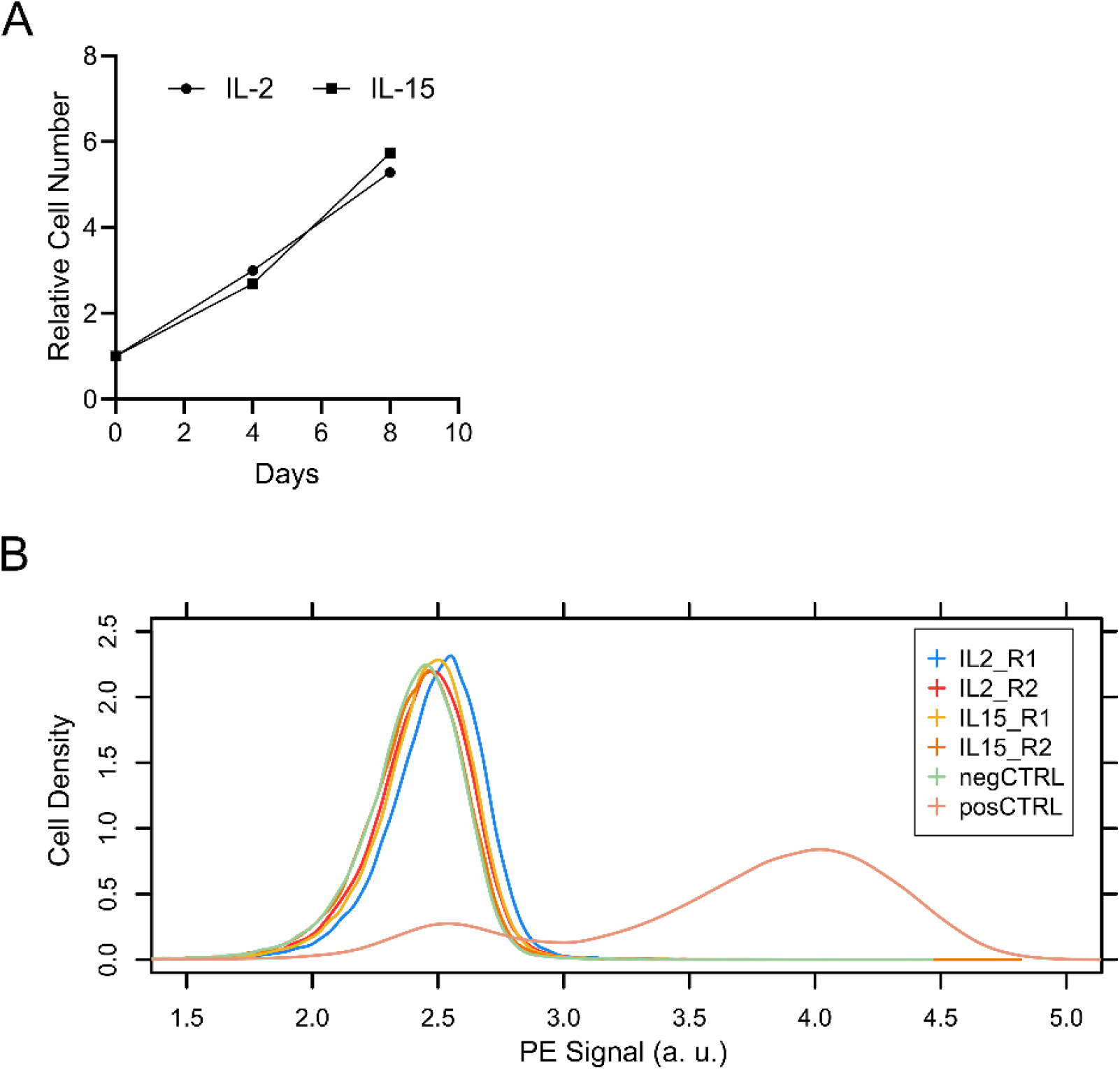
IL-2 and IL-15 induce proliferation but not IFN*γ* production in NK-92 cells. A) NK-92 cells were grown in media containing either IL-2 or IL-15 at a concentration of 100 U/mL. Cells were counted every fourth day by trypan blue exclusion, and the cell number was normalized to the initial cell number. B) IFNγ production was measured by flow cytometry following starved NK cell treatment with either IL-2 or IL-15 in biological replicate. Treatment with PMA and ionomycin was used as the positive control (posCTRL) while no stimulation was maintained as the negative control (negCTRL).

**Supplemental Figure 2.**
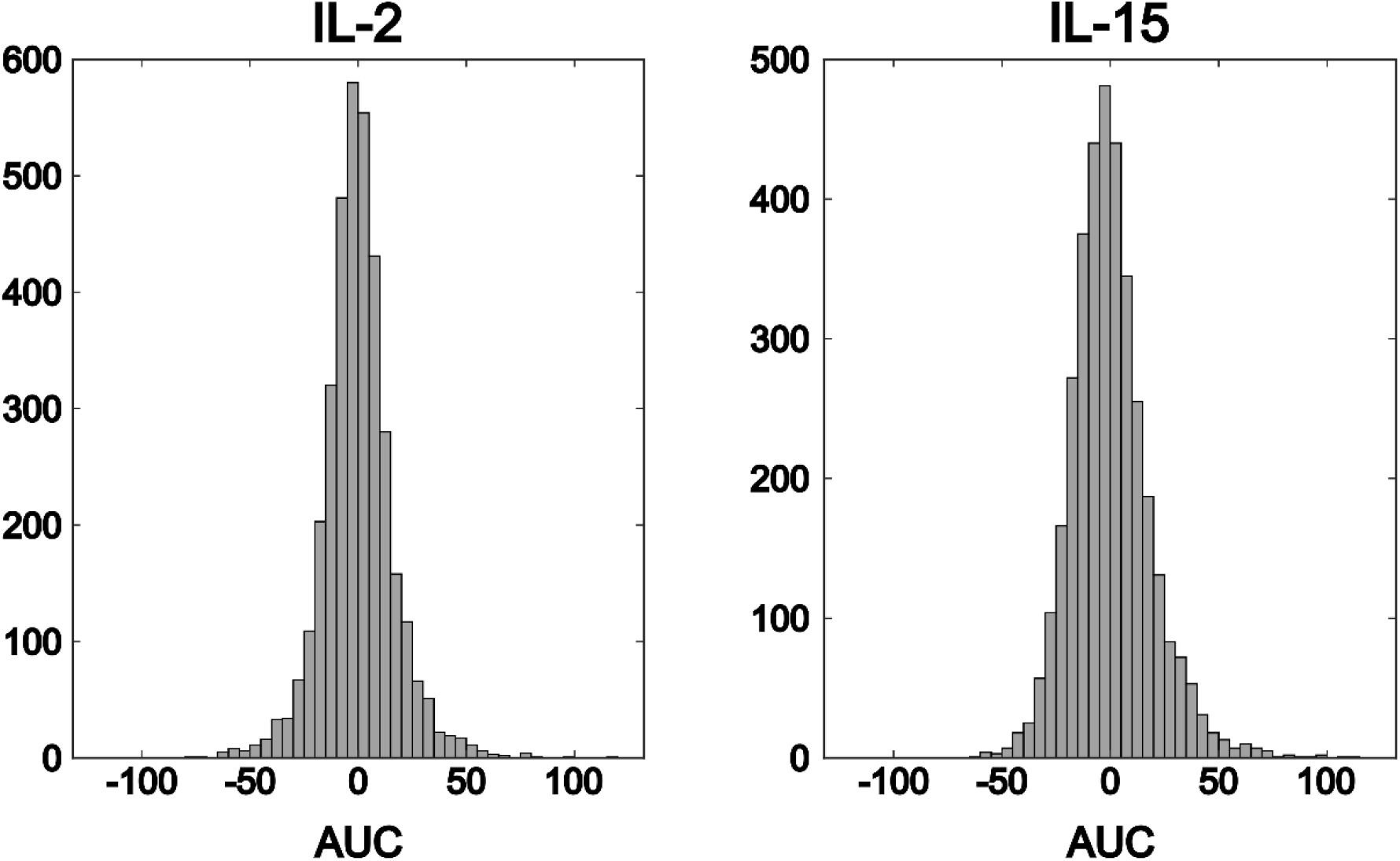
Histograms of IL-2 and IL-15 phospho-proteomic AUC values. For both IL-2 and IL-15, positive and negative AUC values were evenly balanced. For IL-2, mean AUC = -0.1, median AUC = -0.6, and 48.2% of phospho-peptides with AUC > 0. For IL-15, mean AUC = 0.1, median AUC = -1.6, and 46.0% of phospho-peptides with AUC > 0.

